# Extracellular levels of glucose in the hippocampus and striatum during maze training for food or water reward in male rats

**DOI:** 10.1101/2020.04.20.051284

**Authors:** C.J. Scavuzzo, L.A. Newman, P.E. Gold, D.L. Korol

**Author notes:** Corresponding authors: Donna L. Korol, Syracuse University, Department of Biology, Life Sciences Complex, Syracuse, NY 13244, Claire J. Scavuzzo, University of Alberta, Department of Psychology, Edmonton, Alberta, Canada, T6G 2E9.

## Abstract

Peripheral and central administration of glucose potently enhance cognitive functions. The present experiments examined changes in brain extracellular glucose levels while rats were trained to solve hippocampus-sensitive place or striatum-sensitive response learning tasks for food or water reward. During the first minutes of either place or response training, extracellular glucose levels declined in both the hippocampus and striatum, an effect not seen in untrained, rewarded rats. Subsequently, glucose increased in both brain areas under all training conditions, approaching asymptotic levels ∼15-25 min into training. Compared to untrained-food controls, training with food reward resulted in significant glucose increases in the hippocampus but not striatum; striatal glucose levels exhibited large increases to food intake in both trained and untrained groups. In rats trained to find water, glucose levels increased significantly above the values seen in untrained rats in both hippocampus and striatum. In contrast to results seen with lactate measurements, the magnitude of training-related increases in hippocampus and striatum glucose levels did not differ by task under either reward condition. The decreases in glucose early in training might reflect an increase in brain glucose consumption, perhaps triggering increased brain uptake of glucose from blood, as evident in the increases in glucose later in training. Together with past findings measuring lactate levels under the same conditions, the initial decreases in glucose may also stimulate increased production of lactate from astrocytes to support neural metabolism directly and/or to act as a signal to increase blood flow and glucose uptake into the brain.

**Highlights:**

- Glucose levels in hippocampus and striatum decrease at the start of training.
- Glucose levels increase in both brain areas later in training.
- Glucose changes in both brain areas were similar for place and response tasks.
- Glucose levels responded similarly to training for either food and water rewards.
- Early decreases in glucose may trigger increased production astrocytic lactate.

## 1. Introduction

Glucose enhances cognitive functions in rodents as well as in several human populations including healthy young adult and elderly individuals and those with Alzheimer’s disease [1-10]. In past experiments with rats tested on a spatial working memory task, extracellular glucose levels in the hippocampus decreased early and recovered later in testing, the recovery possibly secondary to the onset of testing-related increases in peripheral blood glucose levels [11-14]. In addition to revealing a decrease in extracellular glucose levels in the hippocampus during memory testing, the magnitude of the reduction in extracellular glucose levels was a function of task difficulty, or cognitive load, and also brain structure [11, 12, 15]. Interestingly, striatal extracellular glucose levels failed to decline during spatial working memory and indeed appeared to increase over the course of testing [12]. The significance for learning and memory of these fluxes in extracellular glucose levels is supported by evidence that glucose also enhances learning and memory when microinjected directly into brain targets selected because of their involvement in specific tasks [16-21].

The hippocampus and striatum are two neural systems known to be important participants in selective learning of mazes based on spatial (place) or response (habit) learning, respectively [22-29]. Evidence showing particular importance of the hippocampus and striatum for place and response learning includes double dissociations using lesions or drug disruption of the brain areas [30-34], as well as regulation of the brain areas by direct injections of glutamate, dopamine agonists, glucose and estradiol [21, 35-42]. The hippocampus and striatum also exhibit place and response learning-specific neurophysiological and neurochemical responses [30, 43-51], suggesting selective activation of each structure during the respective canonical task.

One of the mechanisms through which brain glucose might regulate learning and memory is by contributing to astrocytic production and release of lactate into extracellular fluid [2, 3, 14, 52-60]. Recently, we studied the responses of extracellular lactate while rats were trained on place and response mazes [59] that used either food or water as reward. With food reward, extracellular lactate levels increased beyond those of feeding controls during place but not response training, but striatal levels did not exceed the increases in controls for either maze. In contrast, with water reward, lactate did not increase above control levels in the hippocampus during training on either task but increased in the striatum during response but not place learning. Thus, the responses of extracellular lactate to training differed by task but did so with a striking interaction with type of reward.

Because lactate is derived from glucose, largely in astrocytes, the present study examined changes in glucose in the hippocampus and striatum under training and reward conditions matching those of the previous study with lactate [59] to compare and contrast the responses of the two energy substrates to training. In particular, we assessed whether changes in extracellular glucose levels during training would vary in a task by reward manner similar to the results seen with lactate, thereby supporting the view that glucose and lactate are sequentially linked to effects on learning and memory.

## 2. Methods

### 2.1. Subjects

Male Sprague-Dawley rats (3-4 months old; Harlan Laboratories, Indianapolis, IN) were housed individually with free access to food and water under a 12-hour light/dark cycle. All procedures were approved by the University of Illinois Urbana-Champaign and the Syracuse University Institutional Animal Care and Use Committees and are consistent with guidelines detailed in *Guide for Care* and *Use of Laboratory Animals*. The vivarium facilities at both universities are accredited by the Association for Assessment and Accreditation of Laboratory Animal Care.

### 2.2. Cannula Implantation Surgery

Rats were anesthetized using 2-4% isoflurane, injected with 100,000 units of penicillin intramuscularly and either 5 mg/kg Rimadyl or 2.5 mg/kg Flunixin subcutaneously, and placed in a stereotaxic frame while receiving continuous isoflurane though a nose-piece. The skull was exposed through a small incision in the scalp and cleared of blood and fascia. Small burr holes were made in the skull overlying the hippocampus or striatum for guide cannulae (9 mm long, 350 µm diameter, BASi, West Lafayette, IN) and more laterally for hardware to attach the headcap. To permit the later insertion of a glucose biosensors near the time of training, the guide cannulae were placed just dorsal to the hippocampus (AP: -3.8 mm from bregma, ML: + or - 2.5 mm, 0.9 mm ventral to dura) or striatum (AP: 0.2 mm from bregma, ML: + or - 2.0 mm, 1.5 mm ventral to dura, angled 15 degrees away from midline) [59]. The angled approach for the striatum cannulae was needed to accommodate the head stage and later insertion of glucose biosensors [see 59]. Four skull screws were inserted as anchors for dental acrylic used to affix the cannula to the skull and to mount the bottom of the biosensor head stage, which contained a potentiostat, circuitry, and battery for wireless telemetry. Several injections of 0.9% sterile saline (10 ml total, i.p.) were given to the rats post-surgery to provide hydration during recovery. In addition, ibuprofen (Children’s Motrin, 50 mg in 250 ml water bottle) was provided for 24 hrs in the rats’ drinking water for analgesia support after surgery. Rats were allowed to recover for seven days after surgery with free access to food and water and with daily monitoring of health status.

### 2.3. Place and Response Learning

Rats were either food- or water-restricted depending on the reward used during training; water reward was included here to avoid possible confounds of food restriction and reward on brain glucose concentrations. Food-restricted rats received a daily aliquot of standard chow for one week to reduce their body weights to 85% of baseline. During food restriction, rats received several food reward pellets (∼1/3 Frosted Cheerio®) that would be used during training. Water-restricted rats were given water once a day for one week in the same cup that would be used later in the learning tasks. The water restriction schedule reduced the rats’ body weights to 85% of baseline at the end of one week. All rats were handled for 3 min on each of the five days prior to training.

On the training day, each rat was placed in the testing room and allowed to acclimate to the room for approximately one hour. A four-arm radial maze was used with one arm blocked, forming a T-shaped maze as used and detailed in past papers [40, 59, 61-62]. The arms of the maze were 45 cm long ⨯ 14 cm wide ⨯ 7.5 cm high and extended from a 14 cm^2^ center area. For food-rewarded training, each arm had a food cup with perforations at the bottom, permitting placement of food reward below the cup to mitigate the use of odor cues to find the correct arm; the reward pellet in the goal arm sat above the perforated bottom of the cup. For water-rewarded training, the same plastic cups in which the rats had been given water during water restriction were used in the maze and placed in both possible arm choices with only one dish containing water. A single arm choice was allowed for each trial and was determined by passage of all four paws into the chosen arm. Rats were allowed to eat the reward and to drink from the cup during reinforcement for 2-3 seconds before being removed and placed in the holding cage for the intertrial interval of 30 seconds. In both tasks, training continued until rats made 9/10 correct choices or until they reached a maximum of 100 trials. Untrained control rats were fed the food reward every 1.5 minutes or had access to water for 52.5 minutes while in their home cages.

### 2.4. Biosensor measurements of extracellular glucose levels

The biosensor methods were similar to those used before [14, 59, 65]. One day prior to training, rats were briefly anesthetized with 2-4% isoflurane while a glucose biosensor probe (Pinnacle Technology, Inc., Lawrence, KS) was inserted into the guide cannula. The probe was then connected to a wireless potentiostat and the top of the biosensor head stage was attached to enclose the probe system. The rat was returned to its cage overnight to allow the readings of glucose levels to stabilize. The biosensor data were recorded using Pinnacle Technology Laboratory v. 1.6.7 software.

The biosensors extended 3 mm past the end of the guide cannula with the distal 1 mm of the probe coated with glucose oxidase. The current generated by glucose reaction with the enzyme was monitored in 1-sec bins. The responses of extracellular glucose were normalized within rat to the baseline extracellular glucose measures.

After training, the biosensor probes were removed and calibrated to ensure functional integrity. The probes were placed in 8 ml of 0.1 M phosphate-buffered saline until a stable baseline was reached. Every 1.5 min, known amounts of varying concentrations of glucose were then added incrementallyto produce final concentrations of 0.5-7 mM glucose. Resultant changes in current were recorded to create a linear standard curve, which was subsequently used to convert the current recorded during testing to molar concentration of glucose. The biosensors were also coated with ascorbate oxidase to prevent ascorbate interference with the glucose probe; if the coating is compromised, readings reflect both glucose and ascorbic acid.

Therefore, after glucose calibration, ascorbic acid (range: 200-400 µM) was added to the solution and recordings were taken to assess probe integrity. Results from probes that did not have linear responses to glucose, that responded to ascorbic acid, or that malfunctioned during *in vivo* measurements were excluded from analysis (N = 9).

### 2.5. Histology

Upon completion of testing, the rats were anesthetized with 2-4% isoflurane followed by an overdose of pentobarbital (75 mg/kg; Fatal Plus, Vortech Pharmaceuticals, Dearborn, MI). The brain was rapidly extracted and immersed in 10% formalin for at least 2 days, followed by 20% glycerol for 2 days at 4°C. Brains were sectioned in a cryostat and stained with cresyl violet for histological assessments of cannula placements.

### 2.6 Statistical Analyses

Because several rats had trials to criterion scores at the maximum 100 trials allowed, the data for trials to criterion were truncated. Therefore, medians and interquartile ranges were used to display central tendencies for trials to criterion and differences on this learning score were evaluated with non-parametric Mann-Whitney U-tests. For neurochemical analyses, samples were collected from rats every second before, during, and after training and collapsed into 10-second bins. Using these bins, changes in extracellular glucose from baseline were charted across time to portray the full dynamic of glucose response to training. To capture training-induced depletion and elevation of glucose, the averaged values during two epochs are reported: Minutes 1-3 after training, when brain glucose was near its maximum decrease from baseline, and minutes 15-25 during training, when glucose was near its maximum rise during training. Training- and food/water intake-related changes in mean extracellular glucose levels were evaluated within groups with paired sample t-tests comparing baseline values just before training with values at 1-3 min and at 15-25 min during training, with alpha = 0.05. The changes in glucose levels at each epoch after training were also evaluated across groups with two-way ANOVAs to identify main effects of and interactions between brain area and task within each reward condition and additionally by planned paired contrasts. One outlier in the food-untrained group min 15-25 was removed from data analyses (Grubb_4_ = 1.47, p=0.05 critical value). A two-way ANOVA also tested the effects of reward condition (food, water) by brain area (hippocampus, striatum) in rewarded-untrained rats.

## 3. Results

Learning scores depicted as trials to criterion are shown in Figure 1 for rats in the four training-reward conditions: place trained-food, place trained-water, response trained-food, or response trained-water. When trained for either food or water, the number of trials to reach criterion did not differ significantly across place and response training (Food, U_12,12_ = 54.5, p>0.2; Water, U_10,9_ = 50.5, p>0.2).

**Figure 1.**
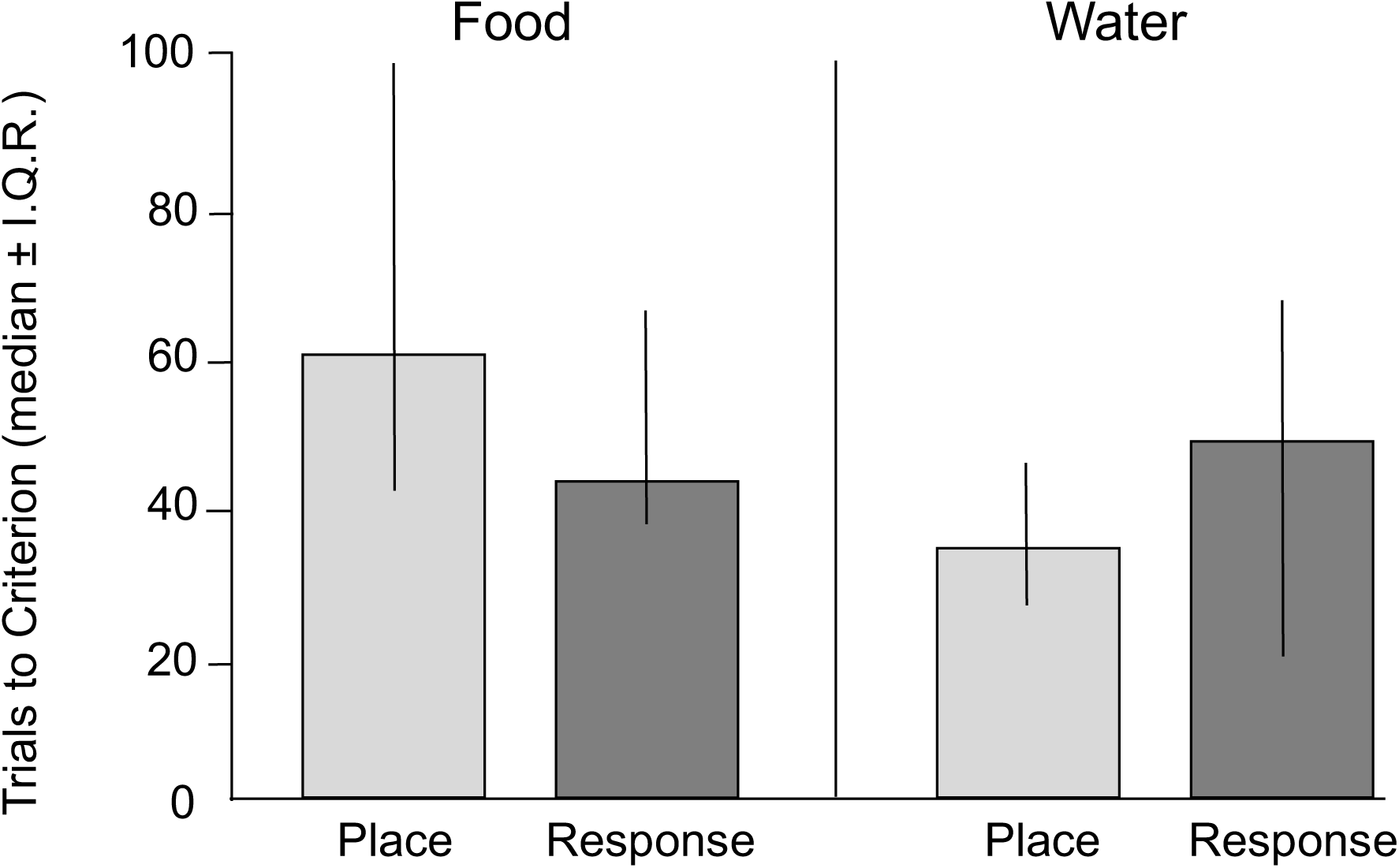
Trials to reach 9/10 correct choices under the four training conditions. Note that rats solved the place and response versions of the maze in similar numbers of trials when trained to find either food or water.

The neurochemical findings revealed two main characteristics of the changes in extracellular glucose levels during training. During the first several minutes of training, extracellular glucose levels decreased from baselines in trained rats. After that time, the concentrations increased substantially throughout the rest of training. To show these different portions of the results graphically, Figures 2-5 focus on the first 5 min of training and Figures 6-9 show the results across the first 40 min of training. Although there was variability in the amount of time required to learn for each training condition all rats required at least 40 min to reach criterion.

**Figure 2.**
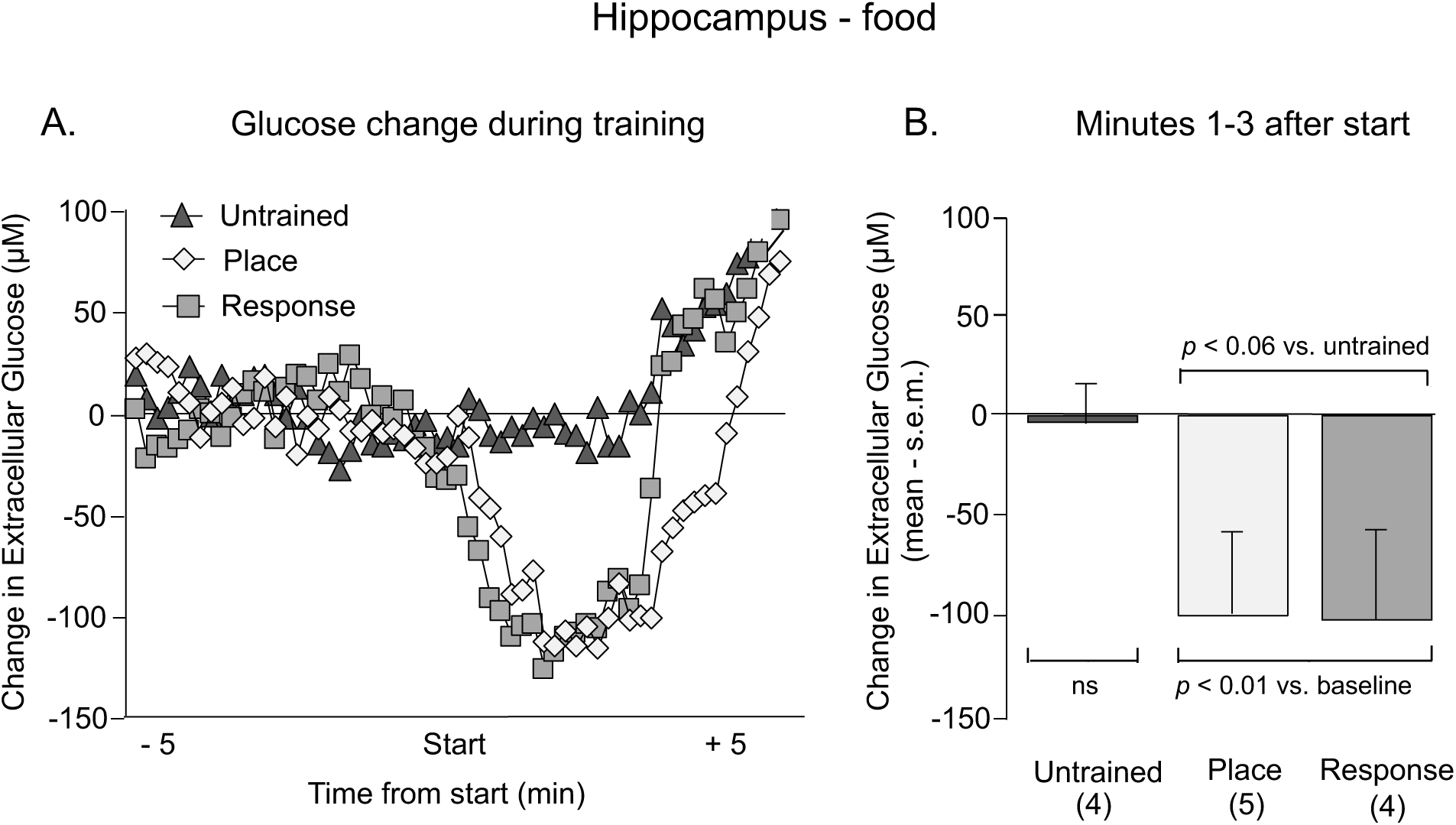
A. Change in extracellular glucose in the hippocampus from 5 min before to 5 min after the start of training for food reward. B. Mean changes in glucose levels during minutes 1-3 after the start of training. Note that the trained groups exhibited significant decreases in glucose levels at this time.

**Figure 3.**
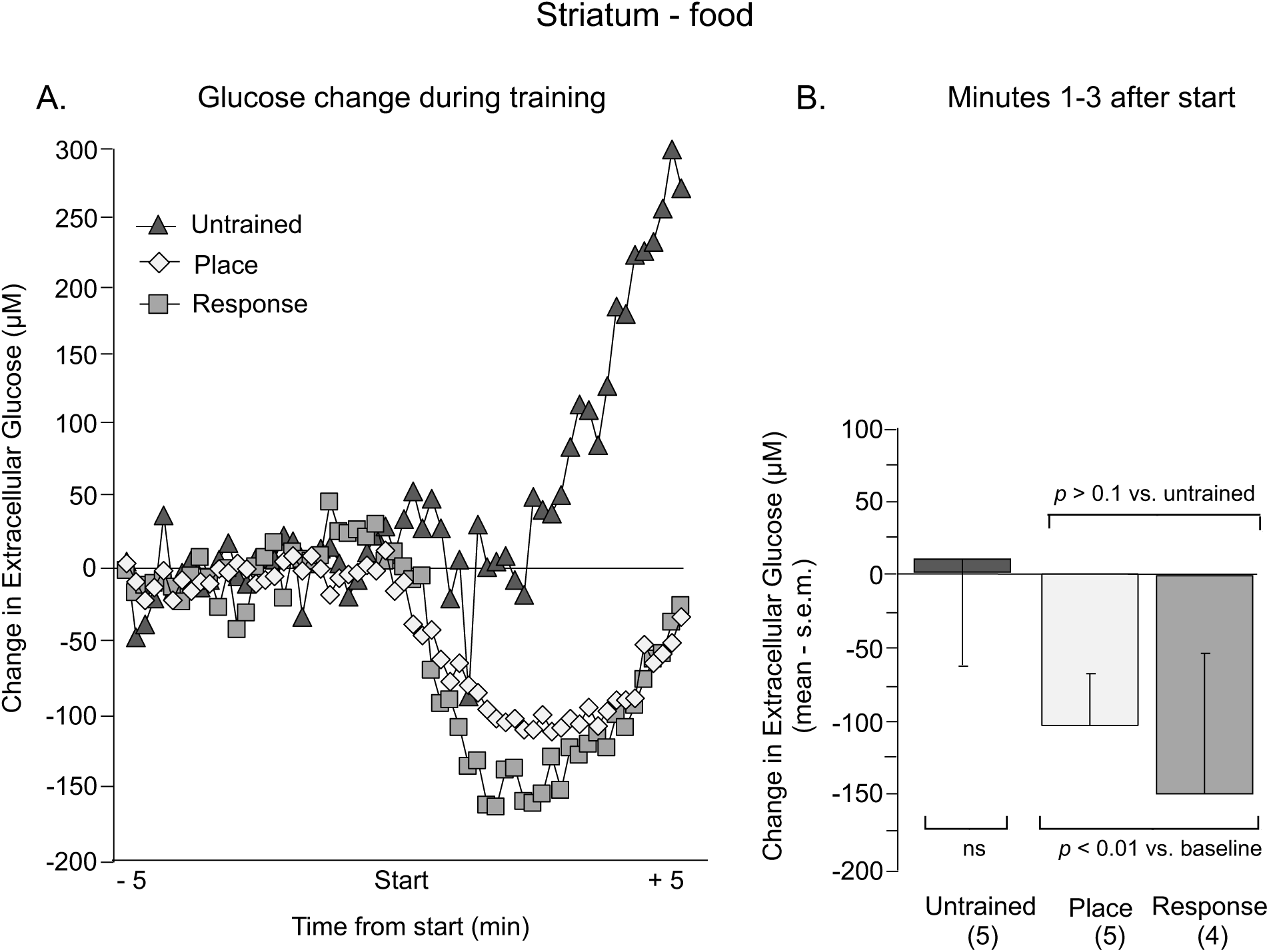
A. Change in extracellular glucose in the striatum from 5 min before to 5 min after the start of training for food reward. B. Mean changes in glucose levels during minutes 1-3 after the start of training. Note that the trained groups exhibited significant decreases in glucose levels at this time.

**Figure 4.**
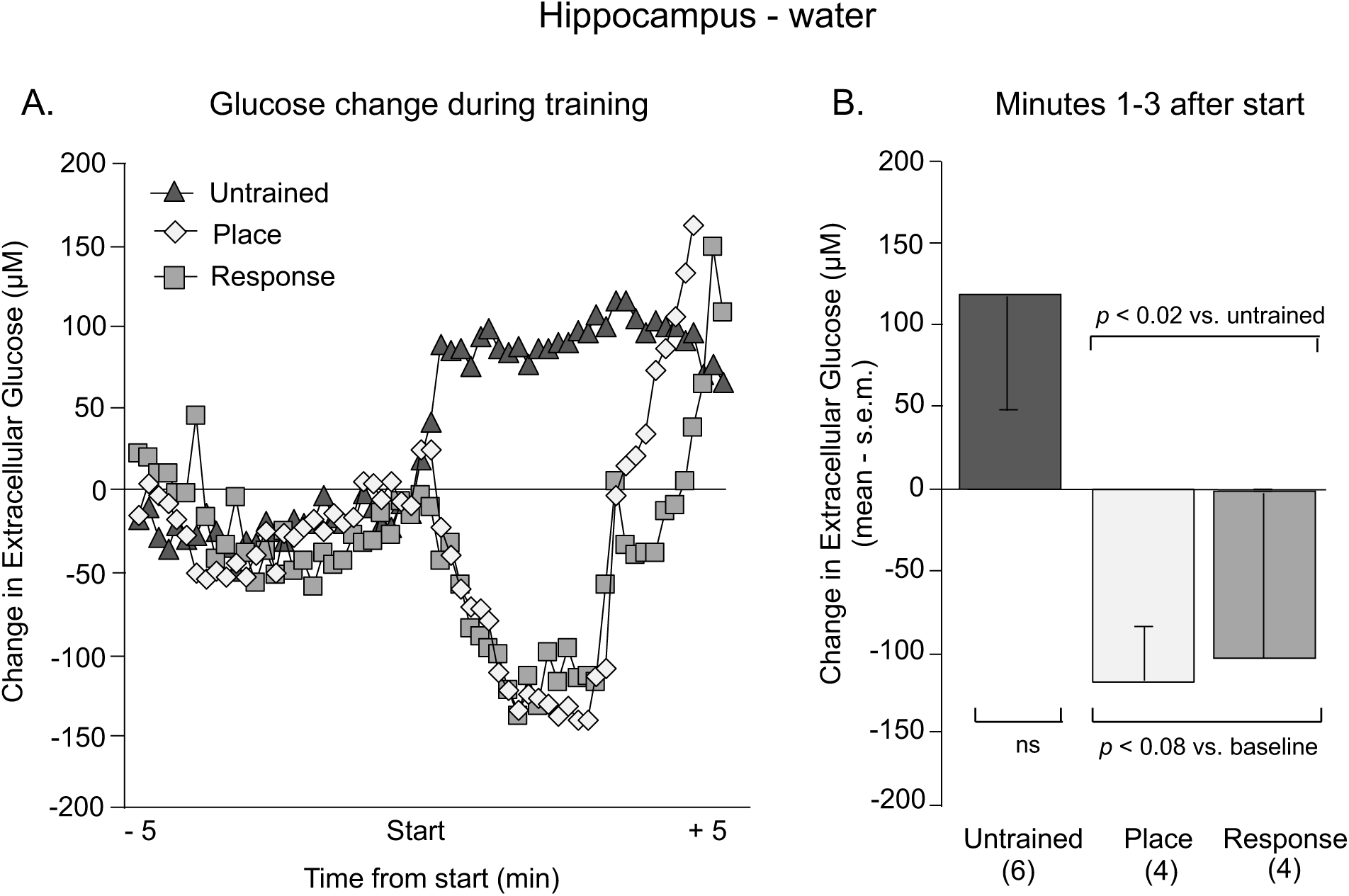
A. Change in extracellular glucose in the hippocampus from 5 min before to 5 min after the start of training for water reward. B. Mean changes in glucose levels during minutes 1-3 after the start of training. Note that the trained groups exhibited decreases in glucose levels at this time. However, this difference was not statistically significant vs. either baseline or vs. untrained groups.

**Figure 5.**
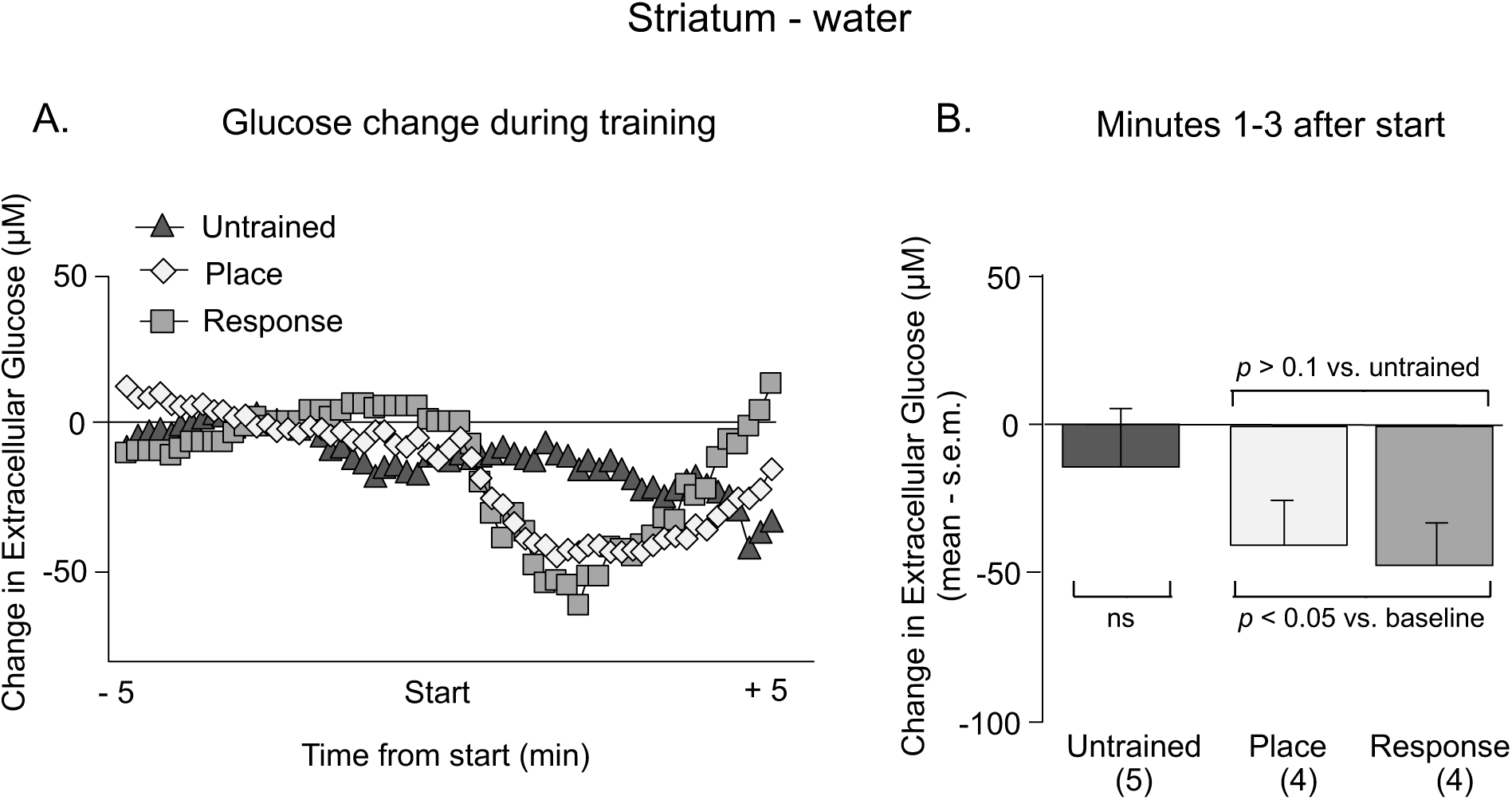
A. Change in extracellular glucose in the striatum from 5 min before to 5 min after the start of training for water reward. B. Mean changes in glucose levels during minutes 1-3 after the start of training. Note that both trained groups but not the untrained group exhibited decreases in glucose levels at this time.

**Figure 6.**
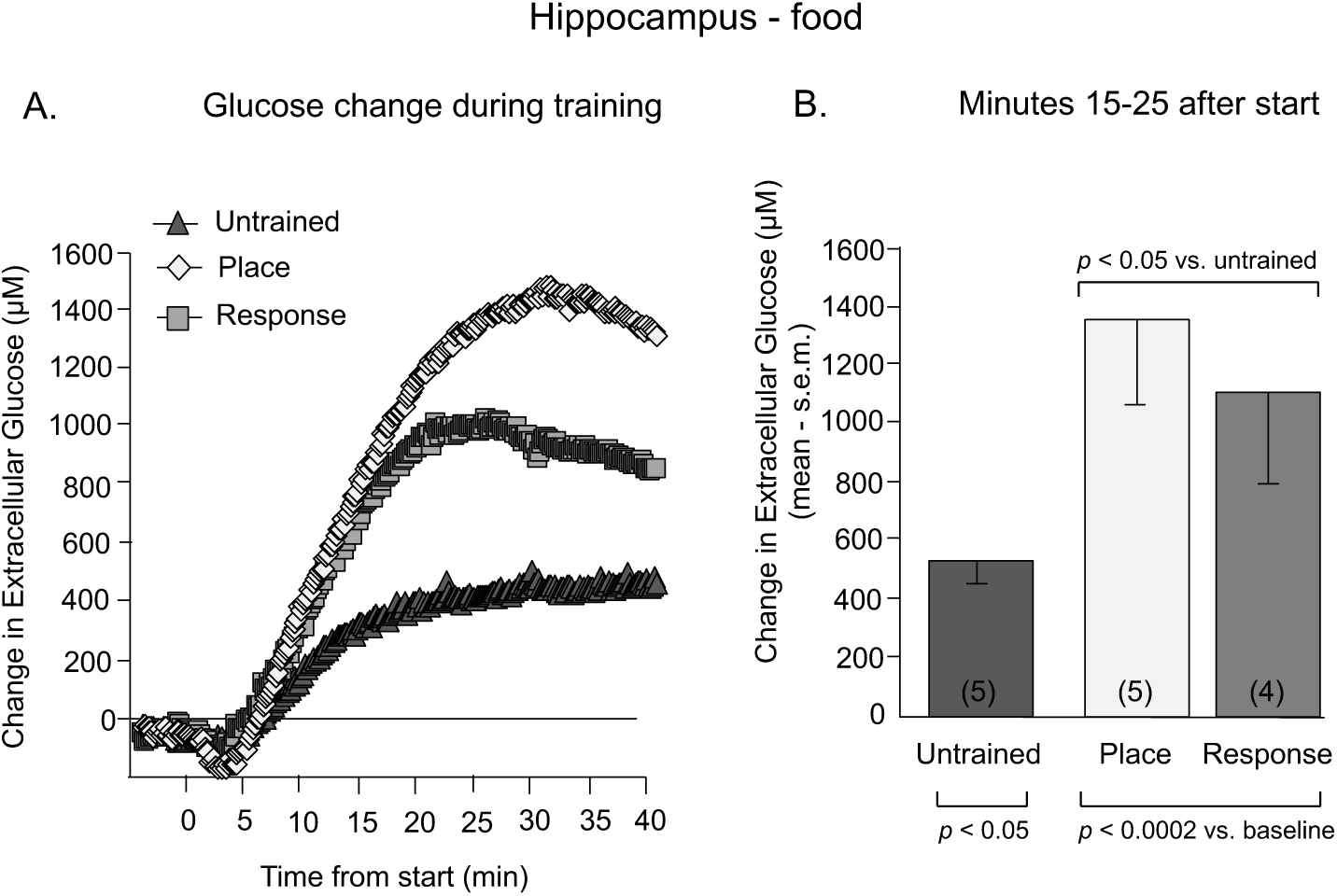
A. Change in extracellular glucose in the hippocampus from 5 min before to 40 min after the start of training for food reward. B. Mean changes in glucose levels during minutes 15-25 after the start of training. All groups, including the untrained group, exhibited significant increases in hippocampal glucose levels above baseline Note that glucose levels in trained rats were significantly higher than those of untrained rats.

**Figure 7.**
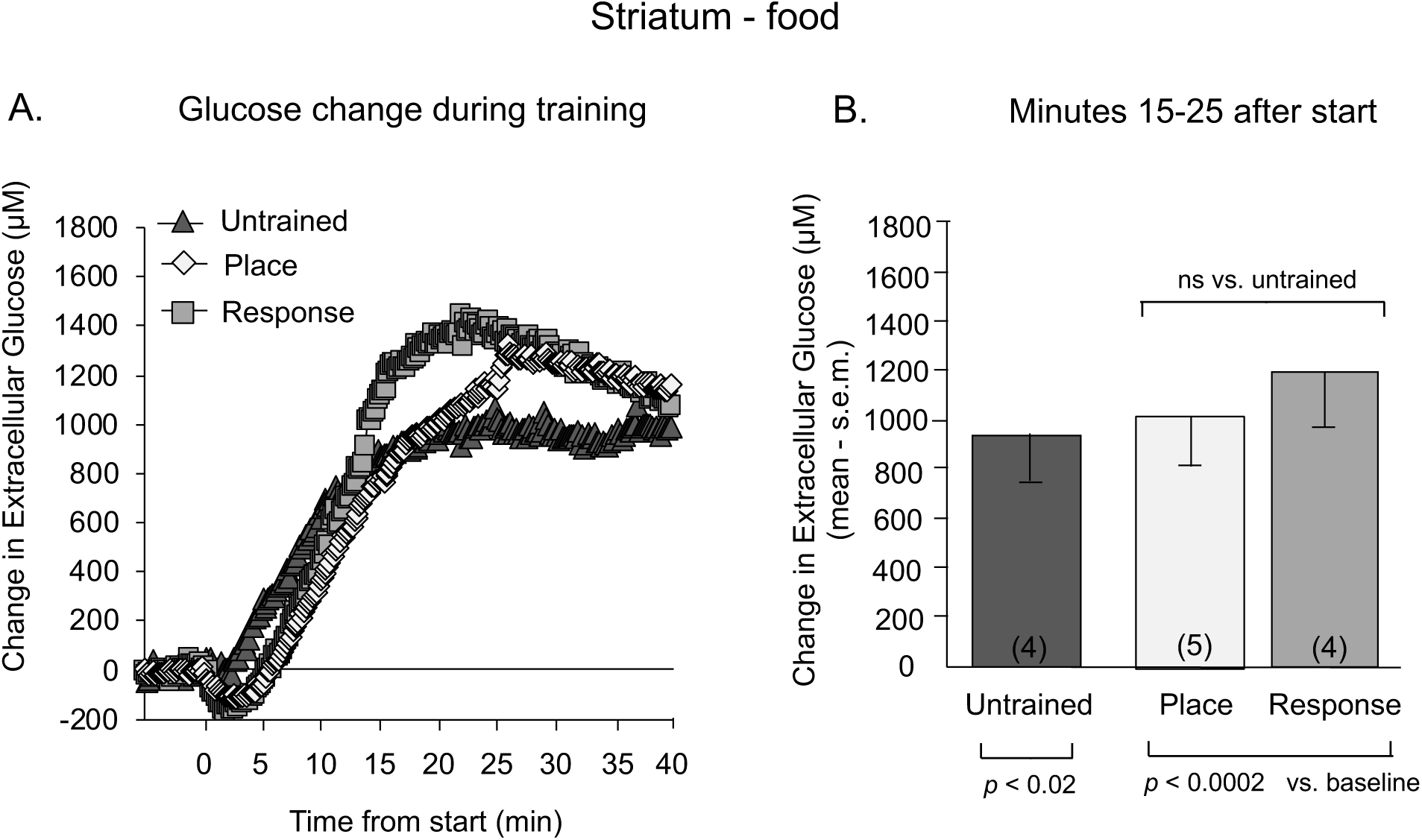
A. Change in extracellular glucose in the striatum from 5 min before to 40 min after the start of training for food reward. B. Mean changes in glucose levels during minutes 15-25 after the start of training. All groups, including the untrained group, exhibited significant ncreases in striatal glucose levels above baseline. Note that glucose levels in trained rats were only slightly higher than those of untrained rats; the difference were not statistically significant.

**Figure 8.**
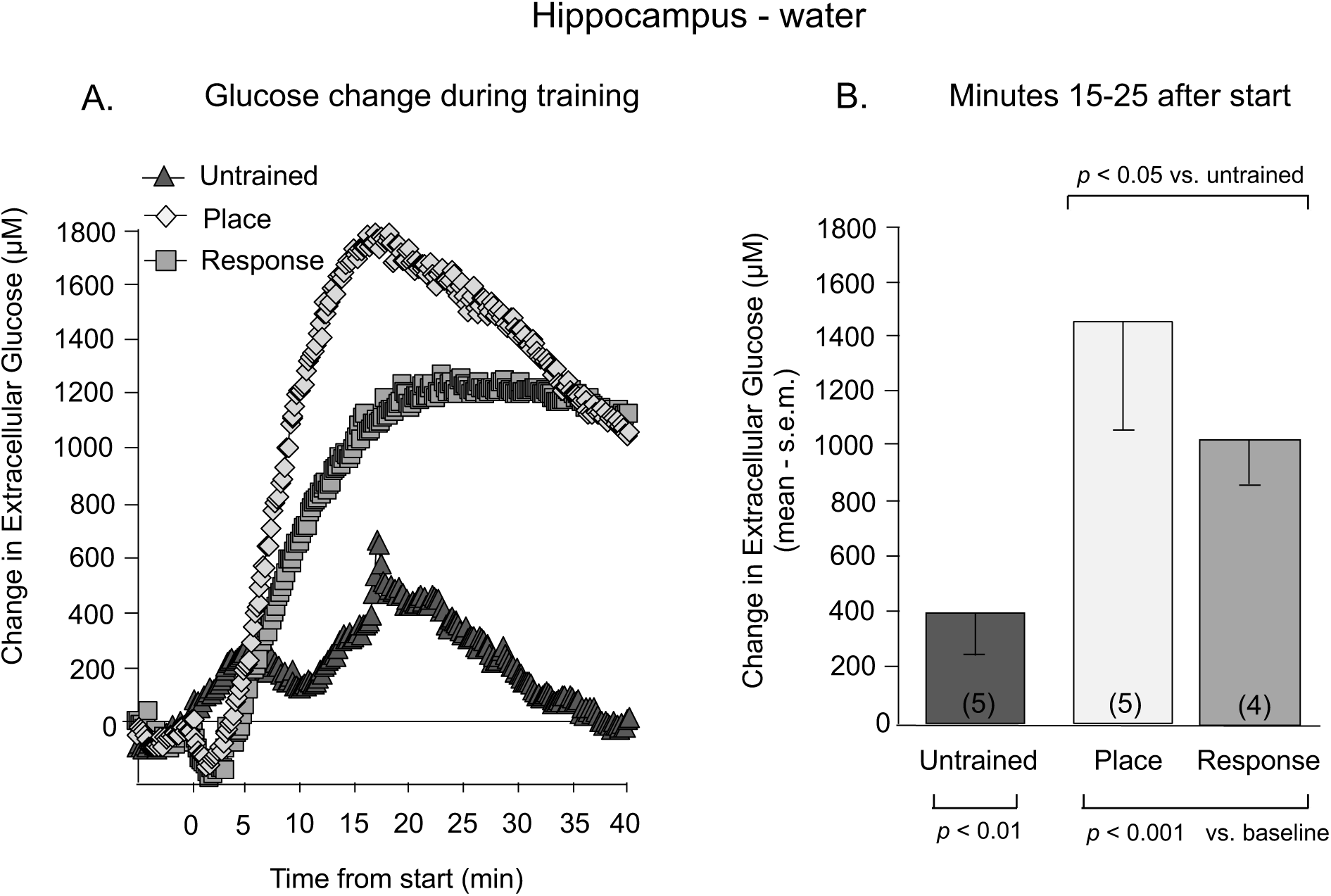
A. Change in extracellular glucose in the hippocampus from 5 min before to 40 min after the start of training for water reward. B. Mean changes in glucose levels during minutes 15-25 after the start of training. All groups, including the untrained group, exhibited significant increases in hippocampal glucose levels above baseline. Note that glucose levels in trained rats were significantly higher than those of untrained rats.

**Figure 9.**
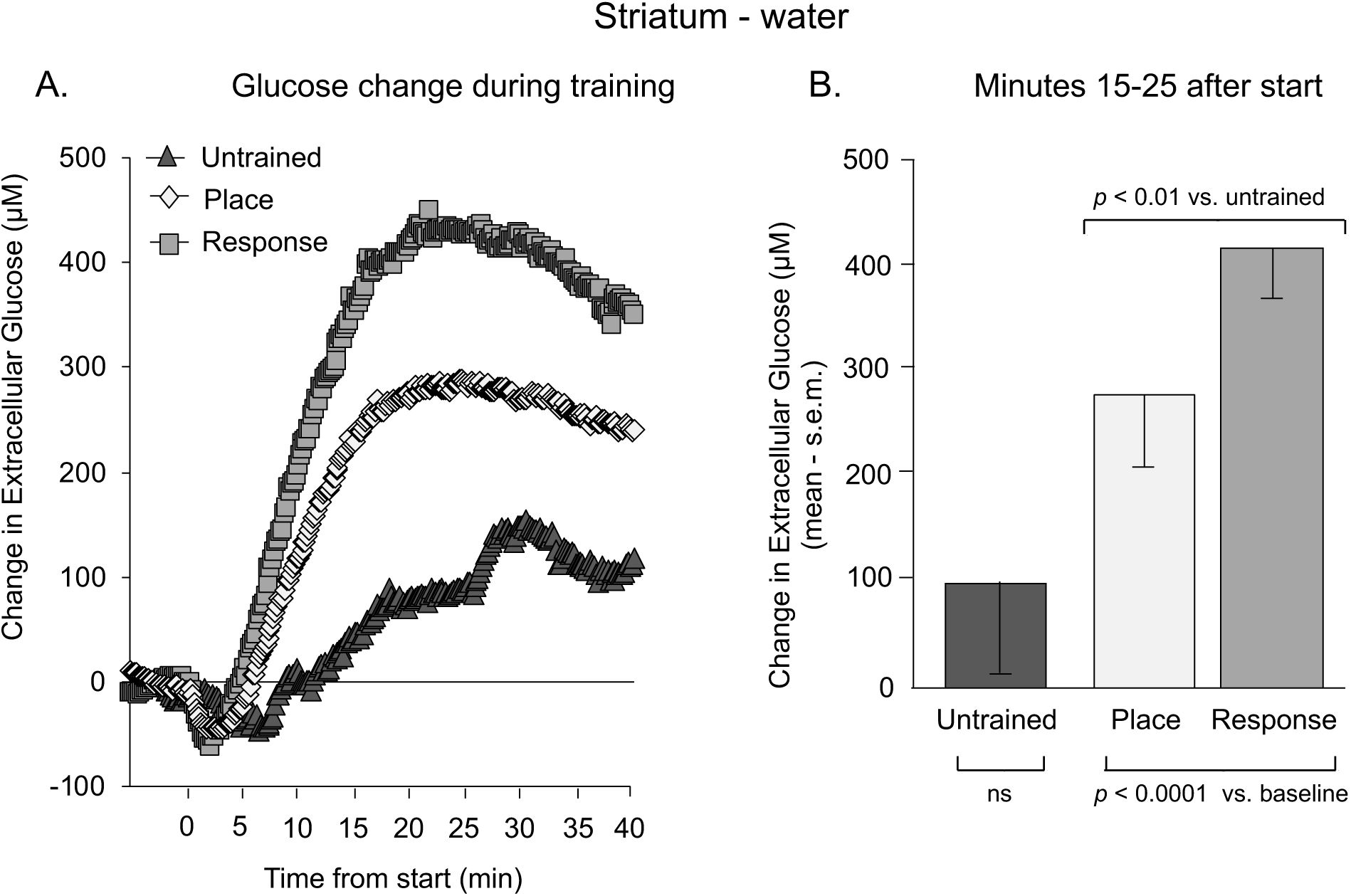
A. Change in extracellular glucose levels in the striatum from 5 min before to 40 min after the start of training for water reward. B. Mean changes in glucose levels during minutes 15-25 after the start of training. The trained groups but not the untrained group had a significant increase in striatal glucose levels above baseline. Note that glucose levels in trained rats were significantly higher than those of untrained rats.

### 3.1. Initial glucose responses to training

The left panels of Figures 2-5 show the responses in 10-sec bins 5 minutes prior to and 5 minutes after the onset of training (or feeding/drinking alone). The right panels show the mean responses during minutes 1-3 of training, i.e. at the time when the responses were near their asymptotic decreases.

#### 3.1.1. Food reward

When rats were trained with food reward (Figures 2 and 3), both hippocampus and striatum demonstrated a rapid decline in ECF glucose at the start of training on both place and response tasks. Evaluation of the double dissociation in the early dip in ECF glucose during training for food revealed no interactions between task and structure (F_1,17_=0.13, p>0.7), with no main effect of structure (F_1,17_=0.09, p > 0.7) or task (F_1,17_=0.19, p > 0.6). Because the early dips in extracellular glucose for food-rewarded rats during place and response training were similar across task in both the hippocampus (Figure 2; t_7_=0.07, p>0.9) and striatum (Figure 3; t_7_=0.47, p>0.6) values from each structure were combined into a ‘trained’ group.

Glucose responses from rats trained for food exhibited hippocampal glucose values that decreased significantly below baseline during minutes 1-3 of training (t_8_=3.57, p<0.01) (Figure 2). In contrast, food-rewarded, untrained rats had hippocampal glucose values that did not significantly change from baseline (t_3_=0.253, p>0.8). The decreases in extracellular glucose levels in food-rewarded trained rats approached significance when compared directly to those of food-rewarded untrained rats (t_11_=2.16, p<0.06). For striatum, food-rewarded trained rats had glucose values during minutes 1-3 (Figure 3) that were again significantly below baseline (t_8_=2.54, p<0.05), while untrained food-rewarded rats had values that did not differ significantly from baseline (t_3_=0.11, p>0.9). There was a trend for greater decreases in striatal glucose levels in food-rewarded trained rats compared to those of food-rewarded untrained rats that was not statistically significant (t_11_=1.46, p>0.1).

#### 3.1.2. Water reward

Like those trained with food rewards, rats rewarded with water (Figures 4 and 5) showed no interaction between brain area and task in extracellular glucose levels during early training (F_1,15_=0.03, p>0.8). There were also no main effects of structure (F_1,15_ = 1.29, p >0.2) or task (F_1,15_= 0.0001, p > 0.9). Because the early decreases in glucose levels between water-rewarded rats receiving place and response training were not significantly different in either the hippocampus (Figure 4; t_6_=0.18, p>0.9) or striatum (Figure 5; t_6_=0.31, p>0.7) those values were combined into ‘trained’ groups within the water-reward condition as was done with training for food above.

Water-trained rats had hippocampal glucose values during minutes 1-3 that dropped below baseline but did not reach statistical significance (t_7_=2.13, p<0.08). However, the early glucose dip in trained rats was significantly different from rats that were untrained given a water reward (t_12_=2.76, p<0.02). Glucose in the hippocampal samples of water-rewarded untrained rats was actually higher, not lower, than baseline; however, this increase was not statistically significant (t_5_=1.77, p>0.1). For striatal samples (Figure 5), glucose levels in the water-rewarded trained rats from minutes 1-3 were significantly below baseline values (t_7_=4.27, p<0.005) and glucose levels in water-rewarded untrained rats were similar to baseline levels (t_5_=0.61, p>0.5). However, the changes in striatal glucose in water-rewarded trained rats were not significantly different from those of untrained rats (t_12_=1.47, p>0.1).

### 3.2. Late glucose responses to training

Figures 6-9 show the changes in extracellular glucose levels in hippocampus and striatum across 40 minutes of training. The left panels show the responses in 10-sec bins during that time. The right panels show the mean responses during minutes 15-25 of training, i.e. a time when the responses are near their asymptotic maxima.

#### 3.2.1. Food reward

When rats were trained for food reward (Figures 6 and 7), there were no significant main effects of brain area (F_1,18_=0.08, p > 0.7), of task (F_1,18_= 0.0001, p > 0.9), or their interaction (F_1,18_=1.1, p>0.3) on the later rise in ECF glucose during minutes 15-25. Although the interaction failed to reach significance, the pattern of effects suggests a dissociation in control of hippocampal ECF glucose during place and response training. For example, mean glucose values were higher in the hippocampus during place than during response training yet values in the striatum were higher during response than during place training. In addition, because the results from food-trained rats receiving place and response training were not significantly different for either hippocampus (t_8_=0.66, p>0.5) or striatum (t_8_=0.89, p>0.4) they were pooled to generate “trained” hippocampus and striatum groups.

For hippocampus samples (Figure 6), the food-rewarded trained rats had glucose values from minutes 15-25 that were significantly higher than baseline (t_8_=6.40, p<0.0002). The untrained food-rewarded rats also had hippocampal values that increased significantly from baseline (t_2_=6.83, p<0.05). The increases in glucose levels in food-rewarded trained rats were greater than those of food-rewarded untrained rats (t_11_=2.36, p<0.05), showing an effect of training on glucose levels beyond that of food intake *per se*. For striatum samples (Figure 7), the food-rewarded trained and untrained rats had glucose values from minutes 15-25 that were significantly above baseline (trained: t_8_=6.40, p<0.0002; untrained: t_3_=5.27, p<0.02). The increases in striatal glucose levels in trained rats did not differ significantly from those of untrained rats receiving food reward(t_11_=0.66, p>0.5), showing no additional effect of training on glucose levels beyond food ingestion. Thus, training with food reward resulted in significant increases beyond those seen in food-untrained rats in the hippocampus but not striatum.

#### 3.2.2. Water reward

When rats were trained for water reward (Figures 8 and 9), mean hippocampal ECF glucose values later in training were higher during place compared to response training while striatal responses were greater during response compared to place training. When tested statistically, the interaction between brain area (hippocampus, striatum) and task (place, response) failed to reach significance (F_1,16_=0.92, p>0.3). In addition, there was a main effect of structure (F_1,16_=12.26, p < 0.005) with hippocampal levels rising higher than striatal levels, but no main effect of task (F_1,1_6=0.63, p > 0.4). Because there were no significant differences with each structure across task, the glucose results of place- and response-trained rats were pooled as above, generating “trained” hippocampus and striatum groups.

Hippocampal glucose levels with water reward rose significantly above baseline in both the trained and untrained conditions (trained: t_8_=5.27, p<0.001; untrained: t_4_=2.56, p<0.01; Figure 8). Of note, trained rats had increases in hippocampal glucose levels that were significantly greater than those of untrained rats (t_12_=2.57, p<0.05). For the striatum, glucose levels increased significantly above baseline in water-rewarded trained rats (t_7_=7.78, p<0.0001) but not in untrained rats (t_5_=1.03, p>0.3; Figure 9). The trained rats had glucose levels higher than those seen in untrained rats (t_12_=3.22, p<0.01). Thus, with water reward, training resulted in increases in both hippocampal and striatal glucose levels significantly above those seen in untrained water-rewarded controls.

### 3.3. Glucose responses to rewards in untrained rats

Untrained rats were given food or water rewards to match the pace of consumption in maze-trained rats. In measurements early in reward provision, there were no significant changes in extracellular glucose in either the hippocampus or striatum. These early times likely preceded increases in blood glucose levels after food (or water) ingestion during initial consumption and therefore were not evident in brain measures [59]. However, at minutes 15-25 after the start of reward consumption in untrained rats, hippocampal and striatal glucose responses to the rewards differed by food vs. water and by brain area. For clarity to compare untrained groups across brain region and reward condition, the changes in glucose levels 15-25 min after the start of reward presentation to untrained groups are replotted from earlier figures and shown in Figure 10. In untrained-rewarded rats, there was a significant interaction of reward by brain area (F_1,17_=11.22, p<0.005), driven mainly by the large differences in the striatal glucose response to food and water. Consumption of food reward increased hippocampal and striatal glucose, as did ingestion of water for hippocampal glucose (p’s<0.05); however, there was no change in striatal extracellular glucose in water-rewarded rats (p>0.2). Of note, the increase in striatal glucose levels after water reward was significantly lower than that of each of the other three groups (p’s<0.02), whereas the magnitude of changes in glucose levels in the other groups did not differ significantly from each other (p’s>0.1).

**Figure 10.**
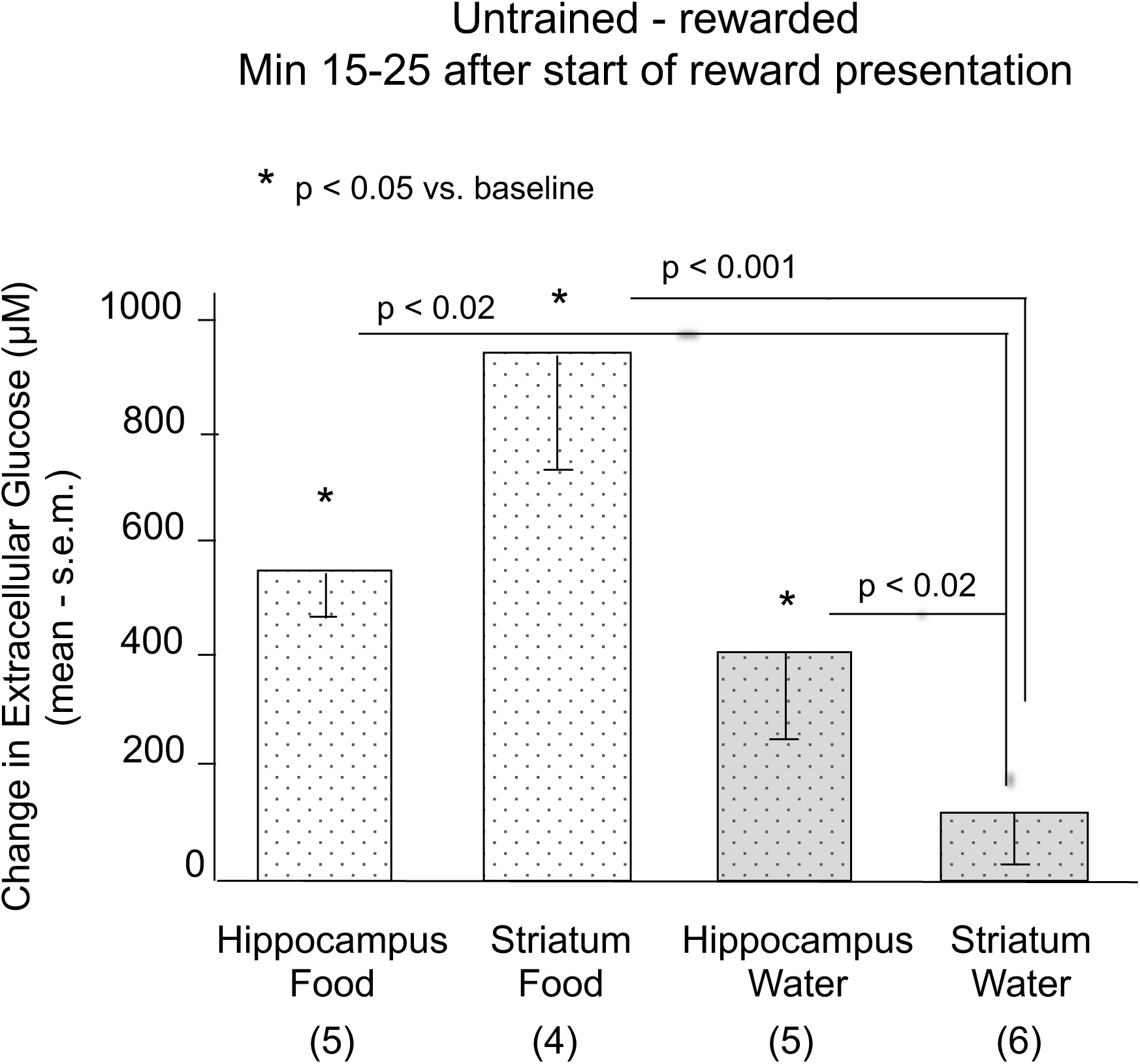
Increases in extracellular glucose levels in the hippocampus and striatum 15-25 min after the start of reward presentation. Food rewards significantly increased extracellular glucose levels in both brain areas. Water rewards significantly increased extracellular glucose levels in the hippocampus but not striatum. In the hippocampus, the increases after food or water did not differ significantly. However, in the striatum, the increase to food was significantly above that to water.

In all rats, accurate placements into hippocampus and striatum were histologically confirmed under light microscopic evaluation; see Figure 11 for examples of hippocampus and striatal placements.

**Figure 11.**
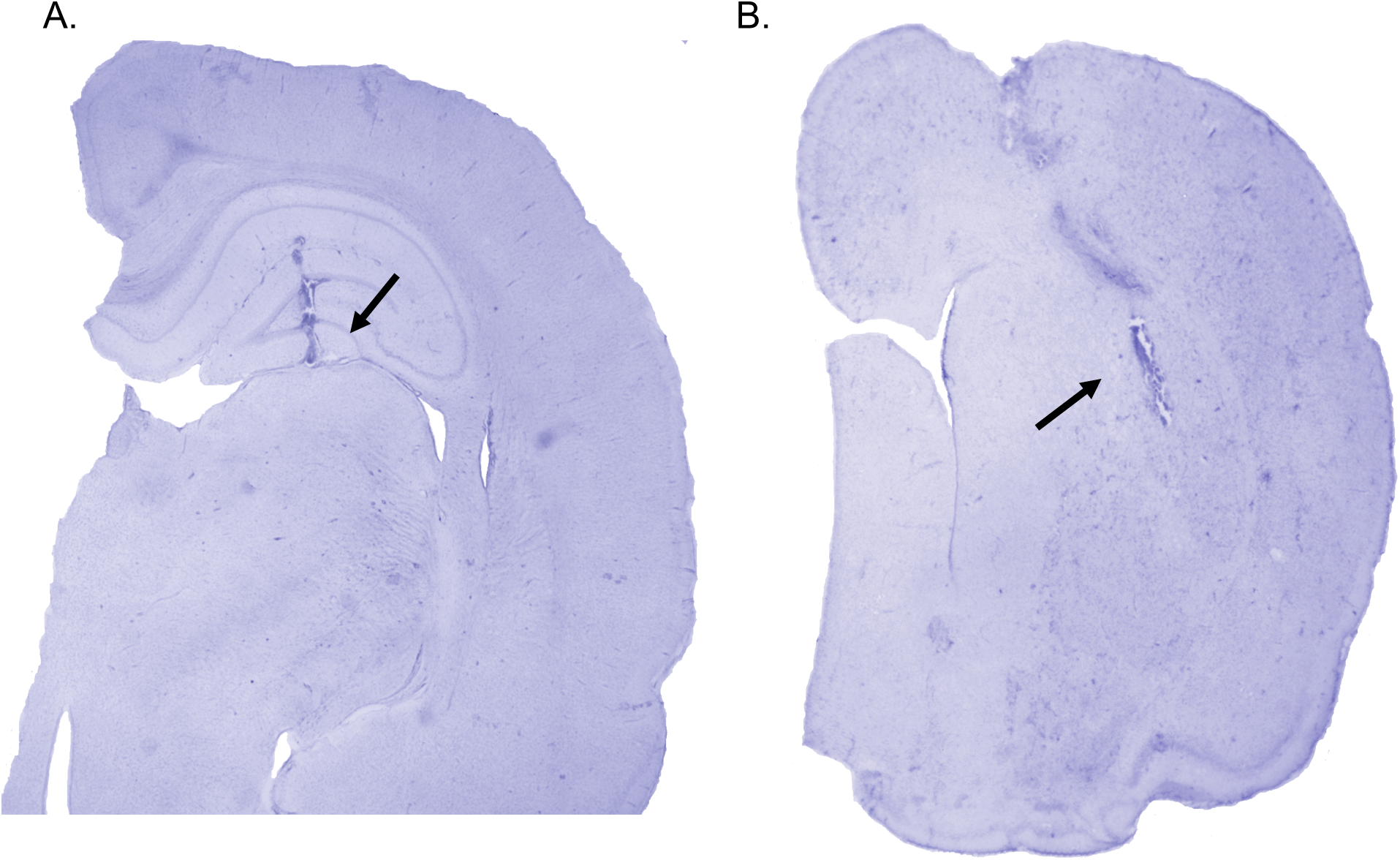
Histological examples of biosensor probe placements in the hippocampus (A) and striatum (B). Probes were located accurately in all rats in this study.

## Discussion

Extracellular glucose levels in the hippocampus and striatum exhibited biphasic responses to training on the place and response versions of the 4-arm radial maze under both food- and water-rewarded conditions. Glucose levels decreased below baseline at the start of training and then increased well above baseline at or around ten minutes into training. An important caveat in interpreting the dynamic extracellular responses of glucose as described in the present report is that the levels evident at any time during behavioral testing reflect momentary equilibria of glucose delivery and its consumption. The methods used here do not distinguish the movement of glucose into or out of different cell types or movement into brain from blood.

The early decreases seen in all trained groups are consistent with previous results obtained with both microdialysis and biosensor measurements of brain glucose levels when rats are tested for spatial working memory on a non-rewarded spontaneous alternation task [11, 14]. The findings that initial decreases in brain extracellular glucose values are evident in both food-and water-rewarded training conditions suggest that the declines were not limited to a condition of reduced glucose availability. In that case, the decrease in glucose levels might be expected to be greater in rats under food restriction than in those under water restriction. Rather, the decreases likely reflect increased brain utilization of glucose in response to the demands of information processing, i.e. uptake of extracellular glucose into neurons and astrocytes [66-70]. Consistent with this view, simultaneous recordings of activated multiunit activity and biosensor measures of glucose concentrations in neocortex reveal an immediate decrease in extracellular glucose concentrations during neural activation [69].

The initial decline in extracellular glucose levels during training was followed by an increase in glucose levels that reached a maximum rise at ∼15-25 min after the start of training. The rise suggests that brain activation recruits additional glucose into extracellular space when needed to support neural activity important for processing learning and memory. The late increase in hippocampal and striatal glucose levels during training above what results from reward alone was clearly evident in water-rewarded rats and for hippocampus with food reward but did not reach significance in food-rewarded trained rats. We have previously reported in untrained rats that food, but not water, intake increases blood glucose [59], suggesting that food rewards alone may have masked training-related increases in glucose during training. However, the use of water reward makes it clear that glucose levels in both the hippocampus and striatum increase as a consequence of place or response training, rather than increasing as a direct function of reward-induced changes in blood glucose levels. The increase in extracellular glucose levels is consistent with evidence that stimulation of brain areas results in increased blood flow, glucose uptake into neurons and astrocytes, and glycolysis [71-74] and that protein and mRNA levels of cerebral glucose transporter 1 (GLUT1) increase immediately after operant training [75]. GLUT1 expression is tightly correlated with local cerebral glucose utilization [76-78], possibly by regulating brain glucose uptake. The plasticity of GLUT1 may contribute to the increase in glucose levels in extracellular fluid samples seen here in the hippocampus and striatum during place and response training.

Like glucose, lactate is also responsive to training and is Because the present experiment mirrored the design of our previous study [59] of changes in extracellular hippocampal and striatal lactate levels during place and response training, it is possible to compare the time-dependent responses of the two energy substrates. Like glucose in the present report, the earlier experiment showed that lactate levels increased in both the hippocampus and striatum during both place and response training. However, lactate levels exhibited task-related differences in the magnitude of increases in lactate levels in the hippocampus and striatum that depended on the type of reward, i.e. food vs. water. In contrast, in the present experiment neither the decreases in glucose early in training nor the increases in glucose later in training were related to task-dependent changes across brain areas. The absence of dissociations of task by brain area responses of glucose diverge from other dissociations evident when measuring acetylcholine [49], cFos, or cJun [51] in place and response tasks such as those used here. The findings are also different than those seen in dual-solution tasks in which the learning can be based either on place or response strategies. With these training procedures, acetylcholine release [43, 46], pCREB and c-Fos expression [79], Arc expression [80], and histone acetylation [81] each reveal differences across hippocampus and striatum in concert with the canonical strategy used. Thus, the data in the present report suggest that changes in glucose levels are less discriminate than are lactate and other neurochemical measures in terms of marking the neural system predominantly engaged by different training procedures and effective learning strategies. Interestingly, in untrained rats, the striatal glucose response was substantially larger to food versus water, suggesting that striatum may contribute to detecting or distinguishing reward attributes, value, or consequences. The findings also indicate that the relationship between glucose and lactate is complex and not simply inverse or monotonic, but instead varies by task, reward, and brain area, with relative changes in the substrates diverging across conditions.

The different responses of glucose, reported here, and lactate [59] do not readily fit a model directly linking the two energy substrates. That changes in glucose and lactate levels support learning and memory processing is consistent with previous findings showing that direct infusions of either glucose [17-18, 21, 35, 40, 82] or lactate [14, 57, 83] into the brain enhance learning and memory in young and aged rats. Reinforcing the view that these substrates are used to support energy metabolism needed during cognitive processing is evidence that direct infusions of other substrates, e.g. pyruvate or the ketone body, β-hydroxybutyrate (B3HB), at the time of inhibitory avoidance training or spontaneous alternation testing also enhance memory functions [84-85].

A consistent finding across measurements of lactate [59] and glucose reported here is that extracellular levels of both glucose and lactate increase after the first few minutes of training and do so robustly. The finding that training induces large increases in both glucose and lactate, together with the ability of the energy substrates to enhance learning and memory, support the idea that neurons require additional energy substrates to engage in optimal processing of learning and memory information.

In addition to serving as a direct energy substrate for neurons, glucose also enters astrocytes [86] where it can be stored as glycogen. According to the astrocyte-neuron lactate shuttle hypothesis, glycogen can then be broken down to lactate, which is delivered to neurons for energy needed to support neural plasticity, learning, and memory [2-3, 56, 70, 87]. The lactate response has been described as reflecting ‘energy on demand’, i.e. provision of an energy substrate to neurons to supplement glucose availability [54, 56, 68, 95-97]. However, reports vary regarding whether neural activation leads to preferentially increased uptake of glucose into astrocytes [86-90] or into neurons [91-94].

During early training, the net demand for and use of glucose is evident as an initial decrease in extracellular levels. A later increase in extracellular glucose during training supports the neuronal energy needs of brain areas engaged in learning and memory to promote mechanisms of neural plasticity. The decrease in available glucose may trigger an increase in extracellular lactate levels from astrocytic glycogenolysis, with lactate then serving as a supplemental fuel for neurons, with the simultaneous increase in both substrates together fueling neuronal activity [98]. Alternatively, if glucose and not lactate is a preferred energy substrate in neurons, the increase in lactate levels may augment glucose availability to neurons acting by roles beyond directly support of the metabolic needs of neurons [99-106]. For example, lactate actions include support for the energetic needs of astrocytes, perhaps thereby sparing glucose consumption by astrocytes in support of glucose use by neurons [102, 106-110]. Lactate also activates GPR81 (or HCAR1) receptors, which can regulate neural excitability [111-112] and possibly modulate neural plasticity and memory [113]. GPR81 receptors are also found on epithelial cells where lactate can serve as a vascular signal [114-117]. In this way, lactate might coordinate region-specific regulation of neurovascular coupling to increase glucose availability in a brain area by task manner. As such, lactate would serve as a mediator of a feed-forward process to promote increases in glucose availability upon demand [118-119]. From this perspective, the findings open the possibility that initial use of glucose, shown here as decreased extracellular levels early in training, leads to increased extracellular lactate levels. Lactate, in turn, may act on GPR81 receptors to signal the vasculature to enable increased brain glucose uptake, shown here as increased glucose levels late in training. Interacting in this manner, the increases in both glucose and lactate could support increased neuronal energy capacity needed for cognitive processing.

## Funding

The work was supported by NSF IOS 13-18490, NIDA DA038798, NIA AG057947, and by P30 AG034464 through the Center on Aging and Policy Studies at Syracuse University.

## CRediT authorship contribution statement

**Claire J. Scavuzzo:** Conceptualization, methodology, investigation, formal analysis, writing – review and editing. **Lori A. Newman:** Conceptualization, methodology, investigation, formal analysis, writing– review and editing. **Paul E. Gold:** Conceptualization, methodology, formal analysis, writing– review and editing, funding acquisition, resources, supervision. **Donna L. Korol:** Conceptualization, methodology, formal analysis, writing, funding acquisition, resources, supervision.

## Declaration of Competing Interests

All authors report they have no conflicts of interest.

## Notes

### Competing Interest Statement

The authors have declared no competing interest.

